# Phosphorylation on PstP regulates cell wall metabolism and antibiotic tolerance in *Mycobacterium smegmatis*

**DOI:** 10.1101/825588

**Authors:** Farah Shamma, Kadamba Papavinasasundaram, Samantha Y. Quintanilla, Aditya Bandekar, Christopher Sassetti, Cara C. Boutte

**Affiliations:** Department of Biology, University of Texas Arlington, Arlington, Texas; Department of Microbiology and Physiological Systems, University of Massachusetts Medical School, Worcester, Massachusetts

**Keywords:** PstP, Serine/Threonine Phosphatase, peptidoglycan metabolism, mycolic acid metabolism, antibiotic tolerance, Mycobacteria, dephosphorylation, starvation, cell wall metabolism, CwlM

## Abstract

*Mycobacterium tuberculosis* and its relatives, like many bacteria, have dynamic cell walls that respond to environmental stresses. Modulation of cell wall metabolism in stress is thought to be responsible for decreased permeability and increased tolerance to antibiotics. The signaling systems that control cell wall metabolism under stress, however, are poorly understood. Here, we examine the cell wall regulatory function of a key cell wall regulator, the Serine Threonine Phosphatase PstP, in the model organism *Mycobacterium smegmatis*. We show that the peptidoglycan regulator CwlM is a substrate of PstP. We find that a phospho-mimetic mutation, *pstP* T171E, slows growth, mis-regulates both mycolic acid and peptidoglycan metabolism in different conditions, and interferes with antibiotic tolerance. These data suggest that phosphorylation on PstP affects its activity against various substrates and is important in the transition between growth and stasis.

**Importance:** Regulation of cell wall assembly is essential for bacterial survival and contributes to pathogenesis and antibiotic tolerance in mycobacteria, including pathogens such as *Mycobacterium tuberculosis*. However, little is known about how the cell wall is regulated in stress. We describe a pathway of cell wall modulation in *Mycobacterium smegmatis* through the only essential Ser/Thr phosphatase, PstP. We showed that phosphorylation on PstP is important in regulating peptidoglycan metabolism in the transition to stasis and mycolic acid metabolism in growth. This regulation also affects antibiotic tolerance in growth and stasis. This work helps us to better understand the phosphorylation-mediated cell wall regulation circuitry in Mycobacteria.

## Introduction

Tuberculosis (TB), an infectious disease caused by the bacterium *Mycobacterium tuberculosis* (*Mtb*) is one of the leading causes of death from infectious diseases (1). The fact that TB treatment requires at least a six month regimen with four antibiotics is partly due to the intrinsic antibiotic tolerance of *Mtb* (2, 3). Stressed *Mtb* cells can achieve a dormant or slow-growing state (4, 5) which exhibits antibiotic tolerance (6), cell wall thickening (7) and altered cell-wall staining (4).

The currently accepted cell wall structure of *Mtb* (8) is composed of three covalently linked layers (9): surrounding the plasma membrane, a peptidoglycan (PG) layer is covalently bound to an arabinogalactan layer. A lipid layer composed of mycolic acids surrounds the arabinogalactan layer, and the inner leaflet of this layer is covalently linked to the arabinogalactan (10). The outer leaflet of the mycolic acid layer contains free mycolic acids, trehalose mycolates and other lipids, glycolipids, glycans and proteins (11). The mycolic acid layer, or mycomembrane, is the outer membrane of mycobacteria and is the major contributor to impermeability of the cell wall (12-14).

In addition to serving as a permeability barrier, regulation of the cell wall likely contributes to antibiotic tolerance, either through further changes in permeability (15), or by changing the activity of antibiotic targets (16). Several studies have observed changes in the cell wall under stress (7, 15, 17, 18). These cell wall changes have been shown to correlate with increased antibiotic tolerance (19–21). This has led the prevalent model that stress-induced regulation of the cell wall contributes to antibiotic tolerance (22). While most of the extant data to support this model is correlative, we recently identified a mutant in *Msmeg* which specifically upregulates peptidoglycan metabolism in starvation and also causes decreased antibiotic tolerance in that condition (23). This shows that there is a causal relationship between cell wall regulation and antibiotic tolerance, at least in limited conditions in *Msmeg*.

Reversible protein phosphorylation is a key regulatory tool used by bacteria for environmental signal transduction to regulate cell growth (24-27). In *Mtb*, Serine/Threonine phosphorylation is important in cell wall regulation (28). *Mtb* has 11 Serine/Threonine Protein Kinases (STPKs) (PknA, PknB and PknD-L) and only one Serine/Threonine protein phosphatase (PstP) (29, 30).

Among the STPKs, PknA and PknB are essential for growth, and phosphorylate substrates many involved in cell growth and division (23, 31-34). Some of these substrates are enzymes whose activity is directly altered by phosphorylation. For example, all the enzymes in the FAS-II system of mycolic acid biosynthesis are inhibited by threonine phosphorylation (35-39). There are also cell wall regulators that are not enzymes, but whose phosphorylation by STPKs affects cell shape and growth. For example, the regulator CwlM, once it is phosphorylated by PknB, activates MurA (23), the first enzyme in PG precursor biosynthesis (40). In the transition to starvation, CwlM is rapidly dephosphorylated in *Msmeg* (23). Mis-regulation of MurA activity increases sensitivity to antibiotics in early starvation (23), implying that phospho-regulation of CwlM promotes antibiotic tolerance. CwlM may also regulate other steps of PG synthesis (41). A recent phospho-proteomic study showed that transcriptional repression of the operon that contains both *pstP* and *pknB* leads to increased phosphorylation of CwlM (42). While the effects of the individual genes were not separated (42), this suggests that PstP could dephosphorylate CwlM.

PstP is essential in *Mtb* and *Msmeg* (43, 44). It is a member of the Protein phosphatase 2C (PP2C) subfamily of metal-dependent protein Serine/Threonine phosphatases (45) which strictly require divalent metal ions for activity (46, 47). PP2C phosphatases are involved in responding to environmental signals, regulating metabolic processes, sporulation, cell growth, division and stress response in a diverse range of prokaryotes and eukaryotes (48-53). PstP_*Mtb*_ shares structural folds and conserved residues with the human PP2Cα (54), which serves as the representative of the PP2C family. PstP_*Mtb*_ has an N-terminal cytoplasmic enzymatic domain, a transmembrane pass and a C-terminal extracellular domain (45).

Many of the proteins known to be dephosphorylated by PstP (35, 45, 55-58) are involved in cell wall metabolism; however, the effects of this activity seem to differ. For example, dephosphorylation of CwlM should decrease PG metabolism in stasis (23). But, dephosphorylation of the FAS-II enzymes (35-37, 59-61) should upregulate lipid metabolism in growth. However, PG and lipid metabolism are expected to be coordinated (22). Therefore, PstP must be able to alter substrate specificity in growth and stasis.

PstP_*Mtb*_ is itself phosphorylated on Threonine residues 137, 141, 174 and 290 (55). We hypothesized that phosphorylation of the threonine residues of PstP might help coordinate activity against different substrates through changes in access to substrates, or through toggling catalytic activity against substrates.

We report here that phospho-ablative and phospho-mimetic mutations at the phospho-site T171 of PstP_*Msmeg*_ (T174 in PstP_*Mtb*_) alter growth rate, cell length, cell wall metabolism and antibiotic tolerance in *Msmeg*. Strains of *Msmeg* with *pstP* T171E alleles grow slowly, are unable to properly downregulate PG metabolism and upregulate antibiotic tolerance in the transition to starvation. We observed that the same mutation has nearly opposite effects on mycolic acid layer metabolism. We also report that PstP_*Mtb*_ dephosphorylates CwlM_*Mtb*_.

## Materials and Methods

### Bacterial strains and culture conditions

All *Mycobacterium smegmatis* mc^2^155 ATCC 700084 cultures were started in 7H9 (Becton, Dickinson, Franklin Lakes, NJ) medium containing 5 g/liter bovine serum albumin (BSA), 2 g/liter dextrose, 0.003 g/liter catalase, 0.85 g/liter NaCl, 0.2% glycerol, and 0.05% Tween 80 and incubated at 37°C until log. phase. Hartmans-de Bont (HdB) minimal medium made as described previously (62) without glycerol was used for starvation assays. Serial dilutions of all CFU counts were plated on LB Lennox agar (Fisher BP1427-2).

*E. coli* Top10, XL1Blue and Dh5α were used for cloning and *E. coli* BL21 Codon Plus was used for protein expression. Antibiotic concentrations for *M. smegmatis* were 25 μg/ml kanamycin. 50 μg/ml hygromycin and 20 μg/ml zeocin. Antibiotic concentrations for *E. coli* were 50 μg/ml kanamycin, 25 μg/ml zeocin, 20 μg/ml chloramphenicol and 140 μg/ml ampicillin.

### Strain construction

The PstP_*Msmeg*_-knockdown strain was made first by creating a merodiploid strain and then by deleting the native *pstP_Msmeg_* gene from its chromosomal location. The merodiploid strain was generated by introducing a constitutively expressing *pstP_Mtb_* gene cloned on an StrR plasmid at the L5 attB integration site. The *pstP_Msmeg_* gene (MSMEG_0033) at the native locus was then deleted by RecET-mediated double stranded recombineering approach using a 1.53 kb loxP-hyg-loxP fragment carrying a 125 bp regions flanking the *pstP_Msmeg_* gene, as described (63). The recombineering substrate was generated by two sequential overlapping PCR of the loxP-hyg-loxP substrate present in the plasmid pKM342. The downstream flanking primer used in the first PCR also carried an optimized mycobacterial ribosome binding site in front of the start codon of MSMEG_0032 to facilitate the expression of the genes present downstream of *pstP_Msmeg_ in* the *Msmeg pstP-pknB* operon.

Deletion of the *pstP_Msmeg_* gene was confirmed by PCR amplification and sequencing of the 5’ and 3’ recombinant junctions, and the absence of an internal wild-type *pstP_Msmeg_* PCR product. The *pstP_Mtb_* allele present at the L5 site was then swapped, as described (64), with a tet-regulatable *pstP_Mtb_* allele (RevTetR-P750-*pstP_Mtb_*-DAS tag-L5-Zeo plasmid). The loxP-flanked *hyg* marker present in the chromosomal locus was then removed by expressing Cre from pCre-sacB-Kan, and the Cre plasmid was subsequently cured from this strain by plating on sucrose. We named this strain CB1175.

Different alleles of *pstP* were attained by swapping the Wild-type (WT) allele at L5 site of CB1175 as described (65). In order to do so, the wild-type (WT) and the phosphoablative alleles of *pstP_Msmeg_* alleles were at first cloned individually into a kanamycin resistant-marked L5 vector pCT94 under a TetO promoter to generate vectors pCB1206-1208 and pCB1210, which would swap out the zeocin-resistance marked vector at the L5 site in CB1175. The strong TetO promoter in the vectors pCB1206-1208 and pCB1210 was swapped with an intermediate strength promoter p766TetON6 (cloned from the vector pCB1030 (pGMCgS-TetON-6 sspB) to generate the L5 vectors pCB1282-85. pCB1285 was used as the parent vector later on to clone in the phosphomimetic *pstP_Msmeg_* alleles under p766TetON6.

These kanamycin resistance-marked vector constructs were then used to swap out the zeocin resistance-marked vector at the L5 site of CB1175 to attain different allelic strains of *pstP_Msmeg_* as described (65).

### Growth Curve assay

At least three biological replicates of different *pstP_Msmeg_* allele strains were grown in 7H9 media up to log. phase. The growth curves were performed in non-treated 96 well plate using plate reader (BioTek Synergy neo2 multi mode reader) in 200μl 7H9 media starting at OD_600_=0.1. Exponential growth equation was used to calculate the doubling times of each strain using the least squared ordinary fit method in GraphPad Prism (Version 7.0d). P values were calculated using two-tailed, unpaired t-tests.

### Cell staining

For staining cells in log. phase, 100μl culture in 7H9 was incubated at 37°C with 1μl of 10mM DMN-Tre for 30 minutes and 1μl of 10mM HADA for 15 minutes. Cells were then pelleted and resuspended in 1x phosphate buffered saline (PBS) supplemented with 0.05% Tween 80 and fixed with 10μl of 16% paraformaldehyde (PFA) for 10 minutes at room temperature. Cells were then washed and resuspended in PBS + Tween 80.

For starvation microscopy, cultures were shaken for 4 hours in HdB media without glycerol at 37°C. 500μl of each culture were pelleted and concentrated to 100μl, then incubated at 37°C with 1μl of 10mM DMN-Tre for 1 hour and 3μl of 10mM HADA for 30 minutes. Cells were then washed and fixed as above. The total time of starvation before fixation was 5.5 hours.

### Microscopy and Image Analysis

Cells were imaged with a Nikon Ti-2 widefield epifluorescence microscope with a Photometrics Prime 95B camera and a Plan Apo 100x, 1.45 NA objective lens. The green fluorescence images for DMN-Tre staining were taken with a 470/40nm excitation filter and a 525/50nm emission filter. Blue fluorescence images for HADA staining were taken using 350/50nm excitation filter and 460/50nm emission filter. All images were captured using NIS Elements software and analyzed using FIJI and MicrobeJ (66). For cell detection in MicrobeJ, appropriate parameters for length, width and area were set. The V-snapping cells were split at the septum so that each daughter cell could be considered as a single cell. Any overlapping cells were excluded from analysis.

Length and mean-intensities of HADA and DMN-Tre signals of 300 cells from each strain (100 cells per replicate) were quantified using MicrobeJ. The values of the mean intensities of 300 cells of each strain are represented in the graph as percentages of the highest mean intensity from all the cells in that experiment. An unpaired, two-tailed t-test was performed on the means of the 300 percentage-intensity values of each strain.

### Western Blots

Cultures were grown in 7H9 to OD_600_=0.8 in 10ml 7H9 media, pelleted and resuspended in 500μL PBS with 1mM PMSF and lysed (MiniBeadBeater-16, Model 607, Biospec). Supernatants from the cell lysate were run on 12% resolving Tris-Glycine gels and then transferred onto PVDF membrane (GE Healthcare). Rabbit α-strep antibody (1:1000, Abcam, ab76949) in TBST buffer with 0.5% milk and goat α-rabbit IgG (H+L) HRP conjugated secondary antibody (1:10,000, ThermoFisher Scientific 31460) in TBST were used to detect PstP-strep. For starvation experiments, cultures were first grown to log. phase, then starved in HdB no glycerol media starting at OD=0.5 for 1.5 hour.

For Western blots of *in vitro* assays, samples were run on 12% SDS gel (Mini-Protean TGX, Biorad, 4561046) and then transferred onto PVDF membrane (GE Healthcare). Mouse α-His antibody (1:1000, Genscript A00186) in TBST buffer with 0.5% BSA and goat α-mouse IgG (H+L) HRP conjugated secondary antibody (1:10,000, Invitrogen A28177) were used to detect His-tagged proteins on the blot. The blots were stripped (Thermo Scientific, 21059) and re-probed with Rabbit α-Phospho-Threonine antibody (1:1000, Cell Signaling #9381) and goat α-rabbit IgG (H+L) HRP conjugated secondary antibody (1:10,000, ThermoFisher Scientific 31460) to detect phosphorylation on the blots.

### Antibiotic assays

For antibiotic assays in log. phase, log. phase cultures were diluted in 7H9 media to OD600= 0.05 before treatment. For starvation assays, cells were grown to OD_600_=0.5, pelleted, washed and resuspended in HdB starvation (with no glycerol and 0.05% Tween) media at OD_600_= 0.3 and incubated at 37°C for a total of 5.5 hours. The cultures were then diluted to OD_600_=0.05 before antibiotic treatment. 8 μg/ml and 45 μg/ml meropenem was used for log. phase and starved cultures, respectively. 10 μg/ml and 90 μg/ml isoniazid was added to log. phase and starved cultures, respectively. 100 μg/ml and 900 μg/ml D-cycloserine was used for log. phase and starved cultures, respectively. 50 μg/ml and 360 μg/ml trimethoprim was added to log. phase and starved cultures, respectively. Samples from the culture were serially diluted and plated on LB agar before and after treatment, and colony forming units were calculated.

### Protein Purification

All the proteins were expressed using *E. coli* BL21 Codon Plus cells. N-terminally his-MBP tagged PknB_*Mtb*_ was expressed and purified as described (67). His-PstP_c_WT_*Mtb*_ (1-300 amino acids of the cytosolic domain, (68)) and His-SUMO-CwlM_*Mtb*_ were both expressed overnight by IPTG induction (1mM and 1.3mM, respectively), purified on Ni-NTA resin (G-Biosciences, #786-940 in 5 ml Bio-Scale™ Mini Cartridges, BioRad #7324661), then dialyzed, concentrated and run over size exclusion resin (GE Biosciences Sephacryl S-200 in HiPrep 26/70 column) to obtain soluble proteins. The buffer for His-SUMO-CwlM_*Mtb*_ was 50mM Tris pH 8, 350mM NaCl, 1mM DTT and 10% glycerol. The buffer for His-PstP_c_WT_*Mtb*_ was 50mM Tris pH 7.5, 350mM NaCl, 1mM DTT and 10% glycerol. 20mM imidazole was added to each buffer for lysis and application to the Ni-NTA column, and 250mM imidazole was added for elution. His-PstP_c_T174E_*Mtb*_ was expressed and purified using the same conditions and buffers used for His-PstP_c_WT_*Mtb*_.

### *In vitro* Dephosphorylation assay

Purified His-SUMO-CwlM_*Mtb*_ was phosphorylated with the purified kinase His-MBP-PknB_*Mtb*_ for 1 hour at room temperature in presence of 0.5mM ATP, 1mM MnCl_2_ and buffer (50mM Tris pH 7.5, 250mM NaCl, and 1mM DTT). The amount of kinase was one-tenth of the amount of substrate in the phosphorylation reaction. To stop the kinase reaction by depleting ATP, 0.434 units of calf intestinal alkaline phosphatase (Quick CIP, NEB, MO525S) per μg of His-SUMO-CwlM_*Mtb*_ was added to the reaction mixture and incubated for 1 hour at 37°C. The reaction mixture was then divided into five parts for the different phosphatase samples and a control with buffer.

Two individually expressed and purified batches of both His-PstP_c_WT_*Mtb*_ and His-PstP_c_T1714E_*Mtb*_ were used as biological replicates to perform the dephosphorylation assay. The reaction was carried out at room temperature for up to 90 minutes in presence of phosphatase buffer (50mM Tris pH 7.5, 10mM MnCl_2_, and 1mM DTT). The amount of phosphatase used was half the amount of His-SUMO-CwlM_*Mtb*_.

The intensities of the α-His and the α-Phospho-Threonine signals on the blots were quantified with FIJI. The intensities of the α-His and the α-Phospho-Threonine signals at each time-point were normalized against the respective antibody-signal intensity at 0m. These relative intensities were used to calculate α-Phospho-Threonine/α-His for each time-point and the values were plotted over time using GraphPad Prism (version 7.0d).

## Results

### Phospho-site T171 of PstP_*Msmeg*_ impacts growth rate

PstP is necessary for cell growth in *Msmeg* (42, 43) and phosphorylation increases PstP_*Mtb*_’s activity against small molecule substrates *in vitro* (42, 43, 55). To see if the phosphorylations on PstP regulate cell growth, we made *Msmeg* strains with either phospho-ablative (T>A) or phospho-mimetic (T>E) alleles (69) at each of the three conserved phosphorylation sites of PstP_*Mtb*_ (42, 43, 55) (Fig. 1A) and performed growth curves. We found that biological replicates of the T134A, T134E, T138A and T138E mutant strains had bi-modal distributions of doubling times (Fig. 1B). Phospho-sites T134 and T138 in PstP_*Mtb*_ map to the flap subdomain (54) (Fig. 1A). This subdomain varies greatly in sequence and structure across different PP2C family members and has been shown to be important in regulating substrate binding, specificity and catalytic activity (54, 70–72). Particularly, T138A and T138E variants of the serine threonine phosphatase tPphA from *Thermosynechococcus elongatus* showed differences in substrate reactivity (70). This suggests that phosphorylation at T134 and T138 could be very important in regulating the normal activity of PstP_*Msmeg*_ in the cell. We think that the inconsistent doubling times of those strains result may result from the formation of suppressor mutants, which we will study in future work.

**FIG 1.**
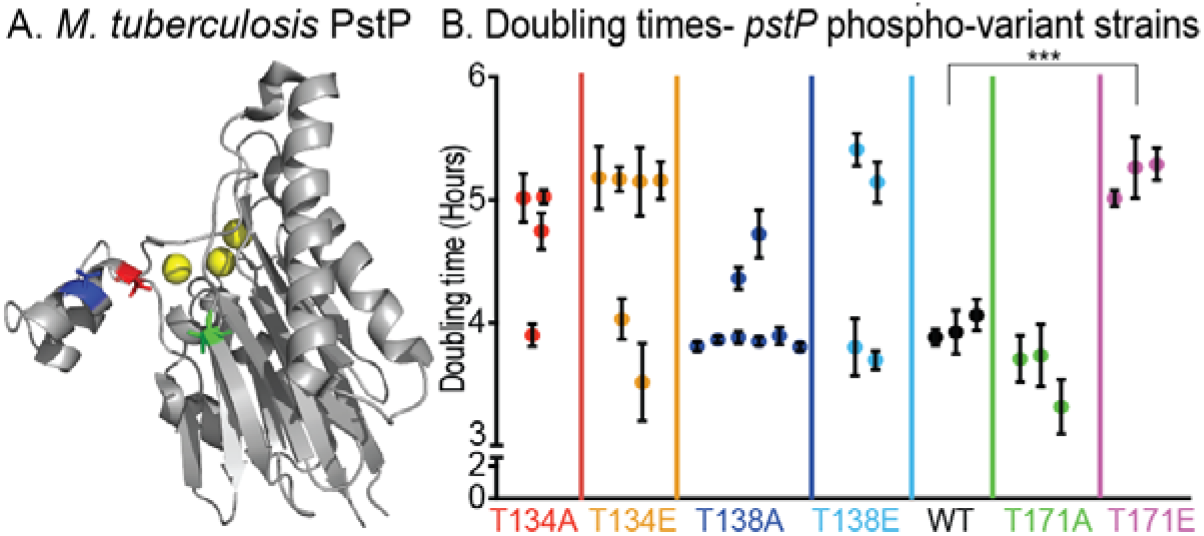
Phospho-site T171 on PstP affects growth. (A) Crystal structure of PstP from *M. tuberculosis (*PstP_*Mtb*_) (54). The threonine (T) sites on PstP_*Mb*_ phosphorylated by the kinases PknA and PknB (55) are highlighted on the structure: red-PstP_*Mb*_ T137 (T134 in PstP_*Msmeg*_), blue-PstP_*Mb*_ T141 (T138 in PstP_*Msmeg*_) and green-PstP_*Mb*_ T174 (T171 in PstP_*Msmeg*_). B) Doubling times of strains containing *pstP_Msmeg_*WT, phospho-ablative mutant alleles *pstP_Msmeg_* T134A, T138A and T171A and phospho-mimetic mutant alleles *pstP_Msmeg_* T134E, T138E and T171E. Each dot is the mean of doubling times from two to three different experiments on different dates of a single isolated clone. The error bars represent the standard deviation. P value= 0.0009.

The *Msmeg* strains with *pstP* T171A and T171E mutations showed consistent and reproducible growth rates (Fig. 1B). The T171A mutants showed no significant difference in doubling time compared to the wild-type, but the T171E grew more slowly than the wild-type. Since T171E mimics constitutive phosphorylation, this result suggests that the continuous presence of a phosphate on T171 downregulates or interferes with cell growth.

### PstP_*Mb*_ WT and PstP_*Mtb*_ T174E dephosphorylate CwlM*_Mtb_ in vitro*

Only a few substrates of PstP have been biochemically verified: some STPKs including PknA and PknB, (45, 55, 57, 58), KasA and KasB (35, 55, 57), and EmbR (56). The STPK PknB phosphorylates CwlM, which is an activator of PG biosynthesis (23). CwlM is rapidly dephosphorylated in the transition to starvation in *Msmeg* (23), and becomes hyper-phosphorylated when PstP is depleted in *Msmeg* (42) which suggests PstP dephosphorylates CwlM.

To test whether PstP and its T174 (T171 in *Msmeg)* phospho-mimetic variant directly dephosphorylate CwlM, we performed an in *vitro* biochemical assay with purified *Mtb* proteins. We purified His-MBP-PknB_*Mtb*_, His-SUMO-CwlM_*Mtb*_ and the cytoplasmic region of PstP_*Mtb*_ that has the catalytic domain (His-PstP_c_WT_*Mtb*_ or PstP_c_T174E_*Mtb*_). PstP dephosphorylates itself rapidly (55), so the purified form is unphosphorylated. We phosphorylated His-SUMO-CwlM_*Mtb*_ by His-MBP-PknB_*Mtb*_, stopped the phosphorylation reaction with Calf Intestinal Phosphatase, and then added His-PstP_c_WT_*Mtb*_ or PstP_c_T174E_*Mtb*_ to His-SUMO-CwlM_*Mtb*_~P. Our control assay with His-SUMO-CwlM~P without PstPcWT_*Mtb*_ or PstPcT174E_*Mtb*_ showed that the phosphorylation on the substrate is stable (Fig. 2A, bottom panel). The phosphorylation signal on His-SUMO-CwlM_*Mtb*_ started decreasing within 5 minutes after addition of His-PstPcWT_*Mtb*_ and kept decreasing over a period of 90 minutes (Fig. 2A, top panel). This is direct biochemical evidence that the PG-regulator CwlM_*Mtb*_ is a substrate of PstP_*Mtb*_. We observed that the WT and T174 phospho-mimetic forms of PstP_*Mtb*_ have no significant differences in activity against His-SUMO-CwlM_*Mtb*_~P *in vitro* (Fig. 2B).

**FIG 2.**
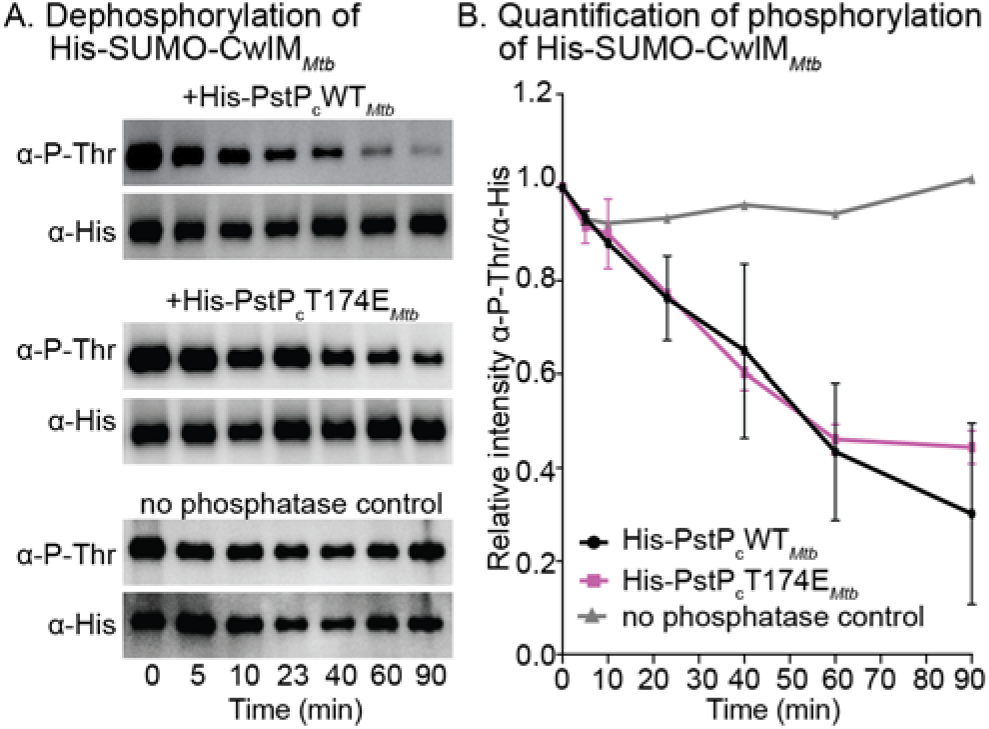
PstP_*Mtb*_ dephosphorylates CwlM_*Mtb*_. (A) α-P-Thr and α-His Western blots of *in vitro* phosphatase reactions with His-PstP_c_WT_*Mtb*_ (top panel) and His-PstP_c_T174E_*Mtb*_ (middle panel), and no phosphatase control (bottom panel) and phosphorylated His-SUMO-CwlM_*Mtb*_. Assay was performed at least twice with two individually purified batches of each phosphatase, one set of images is shown here. (B) Quantification of relative intensities of α-P-Thr over α-His on Western blots. P values were calculated using two-tailed unpaired t-test. All the P values of WT vs T171E at any given time were non-significant. P values of WT vs T171E at 5min = 0.682669, 10min-0.809025, 23min= 0.933929, 40min= 0.831124, 60min= 0.876487 and 90min= 0.545030. The error bars represent standard error of means.

These data show that, *in vitro*, the activity of the catalytic domain of PstP against a single substrate is not affected by a negative charge on T171_*Msmeg*_/T174_*Mtb*_. The phenotypes of the full-length *pstP* T171 phospho-alleles (Fig. 1B, 3, 4 and 5) indicate that this *in vitro* data do not reflect the full complexity of PstP’s regulation *in vivo*.

**FIG 3.**
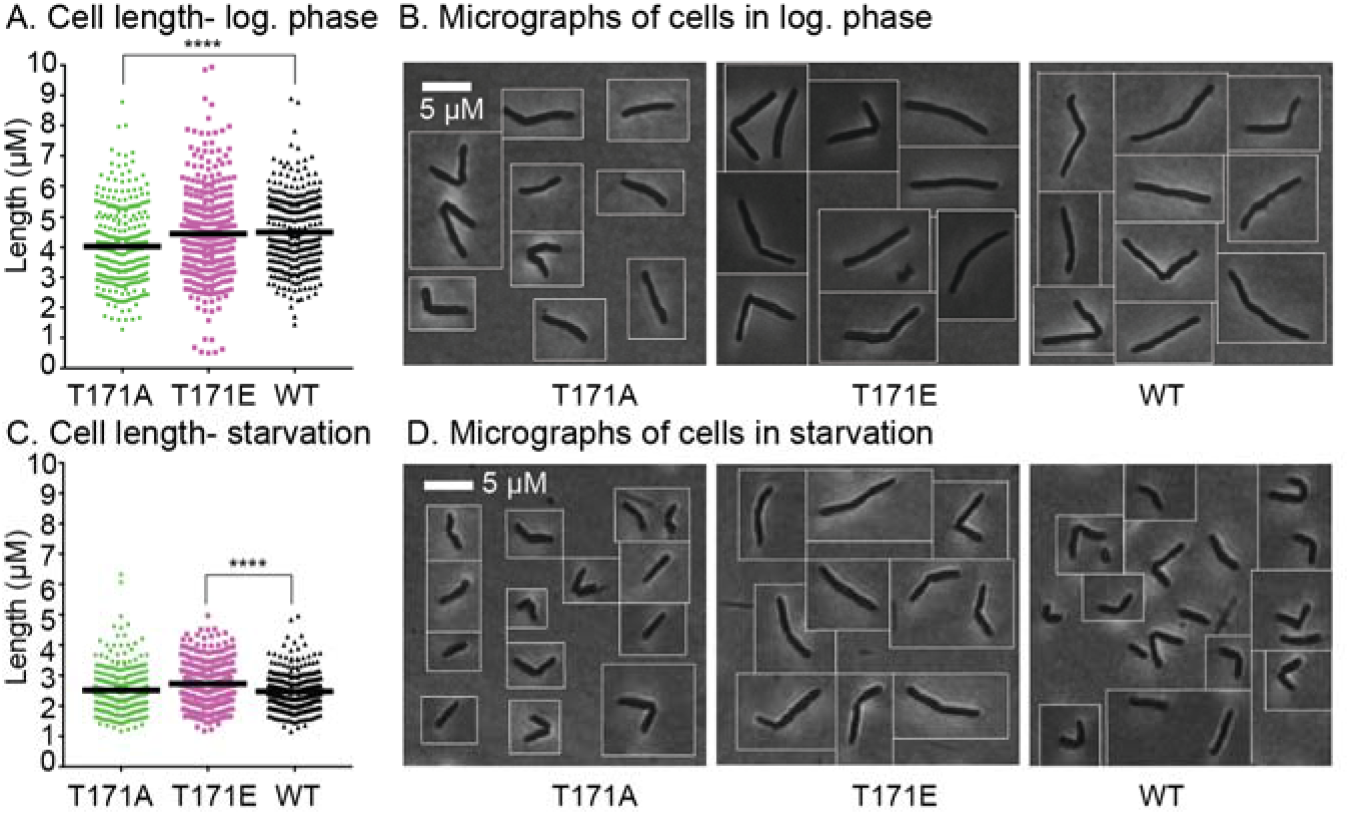
Phospho-site T171 on PstP_*Msmeg*_ is important in regulating cell length. (A) Quantification of cell lengths of isogenic *pstP* allele strains (WT, T17A and T171E) grown in 7H9 in log. phase. 100 cells from each of three biological replicates were measured. P values were calculated by unpaired t-test. P value = 0.000005. (B) Representative phase images of cells from (A). (C) Quantification of cell lengths of isogenic *pstP* allele strains (WT, T17A and T171E) after starvation in HdB with no glycerol for five and a half hours. 100 cells from each of three biological replicates were measured. P values were calculated by unpaired t-test. P value= 0.000003. (D) Representative phase images of cells from (C).

**FIG 4.**
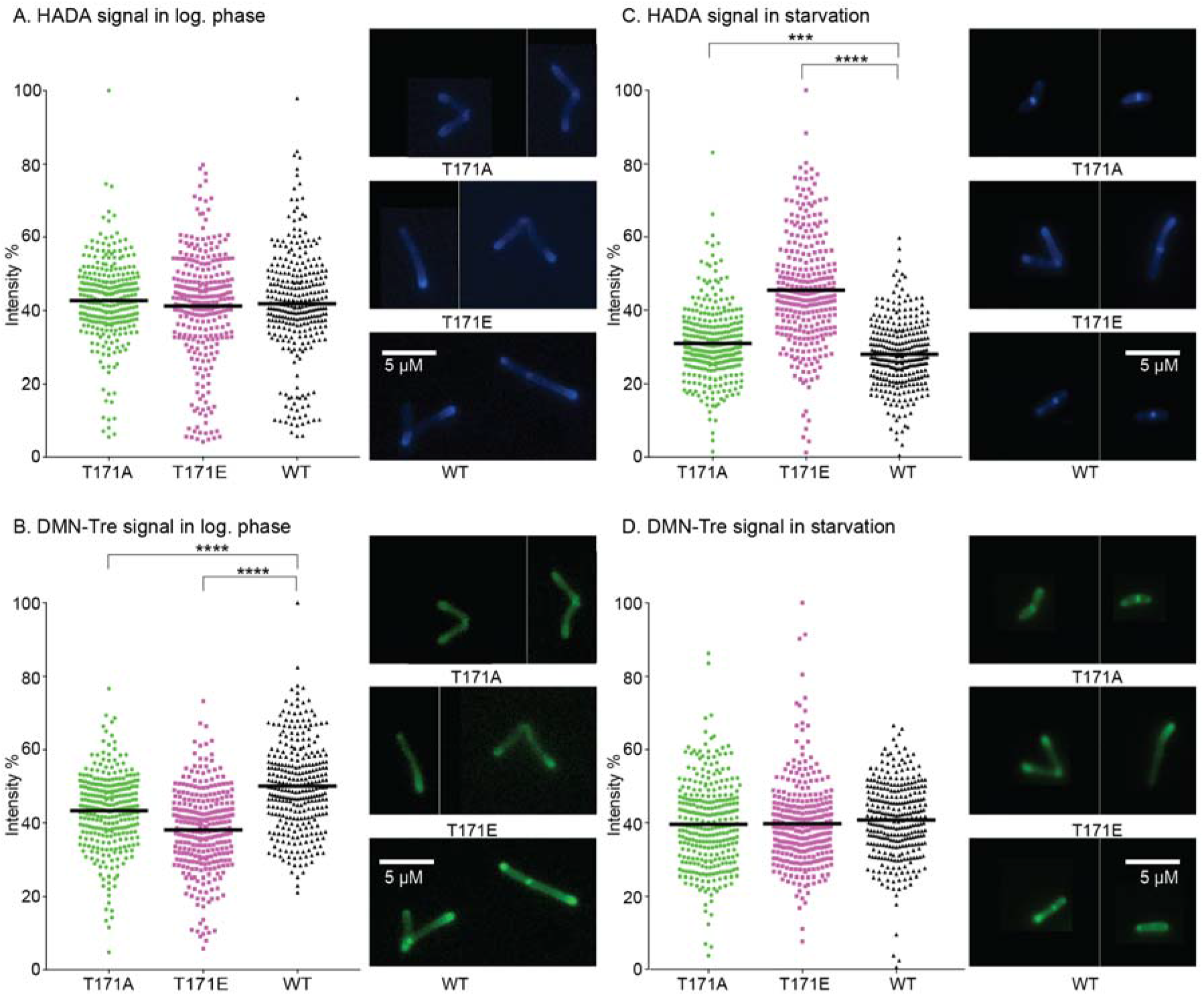
Phospho-site T171 of PstP alters cell wall staining. (A) and (B) Quantification of mean intensities of HADA (A) and DMN-Tre (B) signals of *pstP* allele strains (WT, T17A and T171E) in log. phase cells. P values of both WT vs. T171A and WT vs. T171E= 0.000001. (C) and (D) Quantification of mean intensities of HADA (C) and DMN-Tre (D) signals of starved *pstP* allele strains (WT, T17A and T171E) after 5.5 hours in HdB with no glycerol. P value of WT vs. T171A= 0.0002 and WT vs. T171E = 0.000001.

**FIG 5.**
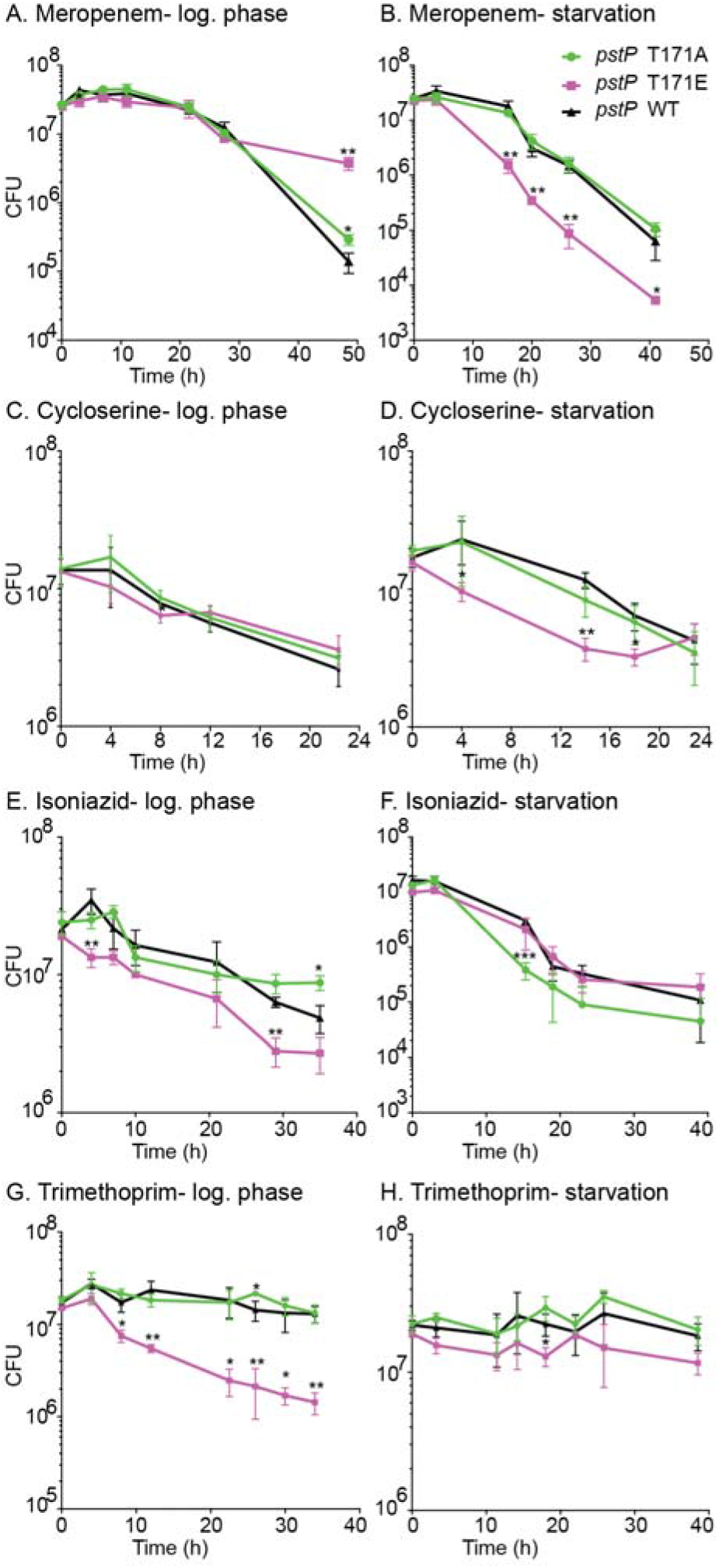
Phospho-site T171 of PstP plays a role in antibiotic sensitivity. Survival of *pstP* allele strains (WT, T17A and T171E) in different media and antibiotics. (A) In 7H9, treated with 8μg/ml of meropenem. P values of WT vs. T171E at 3h=0.043024, WT vs. T171A at 48.5h= 0.001487 and WT vs. T171E at 48.5h = 0.018144. (B) In HdB (no glycerol, 0.05% Tween) for 5.5 hours, then treated with 45μg/ml of meropenem. P values of WT vs. T171E at 16h= 0.002876, 23h=0.006978, 26.3h= 0.003679 and 41h==0.045922. (C) In 7H9, treated with 100μg/ml of D-cycloserine. P values of WT vs. T171E at 8h= 0.032189. (D) In HdB (no Glycerol, 0.05% Tween) for 5.5 hours, then treated with 900μg/ml of D-cycloserine. P values of WT vs. T171E at 3.5h= 0.046062, 14h= 0.001198 and 18h= 0.022088. (E) In 7H9, treated with 10μg/ml of isoniazid. P values of WT vs. T171E at 4h=0.007995, WT vs. T171E at 29h= 0.001978 and WT vs. T171A at 35h= 0.052499. (F) In HdB (no Glycerol, 0.05% Tween) for 5.5 hours, then treated with 90μg/ml of isoniazid. P values of WT vs. T171A at 15.3h=0.000848. (G) In 7H9, treated with 50μg/ml of trimethoprim. P values of WT vs. T171E at 4h= 0.022646, 8h= 0.012762, 12h= 0.005294, 22.5h= 0.014885, 26h= 0.004690, 30h= 0.017293 and 34h= 0.001694. (H) In HdB (no Glycerol, 0.05% Tween) for 5.5 hours, then treated with 360μg/ml of trimethoprim. P values of WT vs. T171A at 18h= 0.023064. All experiments were done at least twice, and representative data are shown. All P values were calculated using two-tailed, unpaired t-test. All error bars represent standard deviation (SD).

### Phospho-site T171 of PstP_*Msmeg*_ regulates cell length

To assess how the phospho-site T171 affects growth, we examined the cell morphology of *pstP* T171 mutants and wild-type (Fig. 3A and B). The quantification of cell length revealed that the *pstP* T171A cells were shorter (mean=4.0261 ± 0.0755) in log. phase than the wild-type cells (mean= 4.5027 ± 0.0704) (Fig. 3A). The *pstP* T171E strain has cell lengths similar to the wild-type (difference between means= 0.05294 ± 0.1163) (Fig. 3A and B) despite the slower growth (difference between means= −1.037 ± 0.1161) (Fig. 1B) in log. phase.

PstP could promote the transition from growth to stasis by downregulating the activity of some cell growth substrates, such as CwlM, PknA or PknB, which all promote growth when phosphorylated (23, 42, 45, 55, 58). PstP likely has dozens of other substrates which may be regulated similarly (23, 42, 45, 55, 58). To test if phospho-site T171 of PstP_*Msmeg*_ affects the transition to stasis, we transferred the strains from log. phase to minimal HdB media with Tween 80 as the only source of carbon, which leads *Msmeg* cells to reductively divide (73). We aerated the cultures for 5.5 hours before imaging (Fig. 3C and D). The effects of phospho-mutations of PstP_*Msmeg*_ on starved cells were the inverse of what we saw in the log. phase. *pstP_Msmeg_* T171E cells in starvation were longer than the wild-type and T171A, and looked like log. phase cells. These data imply that phosphorylation on T171 of PstP_*Msmeg*_ is involved in cell size regulation upon carbon starvation.

### Phospho-site T171 of PstP*_Msmeg_* regulates cell wall metabolism

Since *pstP_Msmeg_* T171 seems to play a role in regulating cell length in growth and stasis, we hypothesized that it affects cell wall metabolism in different phases. To test this, we used fluorescent dyes that preferentially stain metabolically active cell wall(74, 75). We stained T171 allele variant cells from log. phase and after 5.5 hours of carbon starvation with both the fluorescent D-amino acid HADA, which is incorporated into the PG, (74, 76, 77) (Fig. 4A and C, and Fig. S1A and B) and the fluorescent trehalose DMN-Tre, which stains the mycomembrane (75) (Fig. 4B and D, and Fig. S1A and B).

For all experiments, mean intensities of signals from 100 cells from each of three biological replicates of every strain were measured using MicrobeJ. The values of the mean intensities are represented in percentages of the maximum value of all intensities for all strains. P values were calculated by a two-tailed, unpaired t-test.

The PG staining intensity between the strains was the same in log. phase (Fig. 4A). In starvation, the *pstP_Msmeg_* T171E mutant stained much more brightly with HADA than the other strains (Fig. 4C). This suggests that phosphorylation on PstP_*Msmeg*_ T171 may inhibit the downregulation of PG layer biosynthesis in the transition to stasis, but that this phospho-site is not important in modulating PG metabolism during rapid growth.

Staining with DMN-Tre, which correlates with assembly of the trehalose mycolate leaflet of the mycomembrane (75), shows the inverse pattern. The strains stain similarly in starvation (Fig. 4D). In log. phase, however, both mutants show a significant decrease in DMN-Tre signal compared to the wild-type (Fig. 4B), although the *pstP_Msmeg_* T171E mutant has the weakest staining. DMN-Tre is incorporated via Ag85-mediated trehalose mycolate metabolism of the mycomembrane (75). Its fluorescence is sensitive to the hydrophobicity of the membrane and to changes in cytoplasmic mycolic acid metabolism (75).

Our data in Fig. 4A and C suggest that phosphorylation on PstP_*Msmeg*_ T171 impacts PG layer metabolism in starvation, but not growth. But the same phosphorylation appears to regulate the trehalose mycolate metabolism in growth, but not starvation (Fig. 4B and D).

### Phospho-site T171 of PstP_*Msmeg*_ affects antibiotic tolerance

Stresses that arrest cell growth in mycobacteria are associated with increased antibiotic tolerance (15, 18, 78-80). We hypothesized that if *Msmeg* fails to downregulate PG synthesis in starvation, (Fig. 4C), then it might be more susceptible to a PG targeting drug. We treated *pstP_Msmeg_* wild-type, T171A and T171E strains in log. phase and starvation with meropenem, which targets the cross-linking in the PG cell wall (81), and quantified survival by CFU. We saw that the *pstP_Msmeg_* T171E strain was more susceptible in starvation, compared to *pstP_Msmeg_* T171A and wild-type strains (Fig. 5B), but survived similarly in log. phase, except at very late time points when it was more tolerant.

We also treated *pstP_Msmeg_* wild-type, T171A and T171E strains in log. phase and starvation with D-cycloserine, which inhibits incorporation of D-alanine into PG pentapeptides in the cytoplasm (82, 83). The results were similar to those in meropenem: the strains survived similarly in log. phase (Fig. 5C), but the *pstP_Msmeg_* T171E strain was more sensitive in starvation (Fig. 5D). The apparent failure of the *pstP_Msmeg_* T171E strain to downregulate PG synthesis (Fig. 4C) likely makes it more sensitive to both the PG inhibitors in starvation (Fig. 5B and D).

Next, we treated our wild-type and *pstP_Msmeg_* T171 mutant strains with isoniazid, which targets InhA in the FAS-II pathway of mycolic acid synthesis (84). We do not see significant differences in isoniazid sensitivity between the strains in starvation (Fig. 5F). In log. phase, we see that the *pstP_Msmeg_* T171E strain is more susceptible to isoniazid than the *pstP_Msmeg_* T171A and the wild-type strains (Fig. 5E). Our data (Fig. 5E) suggest that phosphorylation on PstP_*Msmeg*_ T171 mis-regulates the mycolic acid biosynthesis pathway of mycomembrane metabolism (Fig. 4C), thus increasing isoniazid susceptibility.

To see if the PstP T171 phospho-site affects susceptibility to a drug that does not target the cell wall, we treated the strains with trimethoprim, which targets thymidine biosynthesis in the cytoplasm (85). We see that, in log. phase, *pstP* T171E is very susceptible to this drug (Fig. 5G), while the wild-type and T171A strains are tolerant. All strains were tolerant to trimethoprim in starvation (Fig. 5H). This shows that mis-regulation of PstP may impact processes beyond cell wall metabolism affecting antibiotic tolerance in turn.

Trimethoprim is a hydrophobic drug which is taken up via passive diffusion (86). Therefore, permeability to trimethoprim is expected to be affected by changes in mycomembrane metabolism in log. phase (Fig. 4C and 5E). It is notable that *pstP* phospho-allele strains in starvation do not exhibit differences in DMN-Tre staining (Fig. 4D) or isoniazid (Fig. 5F) or trimethoprim sensitivity (Fig. 5H), which suggests that susceptibility to trimethoprim, could be determined largely by permeability of the mycomembrane layer. D-cycloserine, on the other hand, is hydrophilic and therefore its uptake is likely dependent on porins (87, 88), and therefore less sensitive to changes in the mycomembrane. So, sensitivity to D-cycloserine (Fig. 5C and D) appears to be largely dependent on regulation of PG metabolism (Fig. 4A and C).

Our data show that phospho-site T171 of PstP regulates mycolic acid layer biosynthesis in growth, and PG layer metabolism in starvation. Mis-regulation of PstP can increase sensitivity to cell wall targeting drugs in both growth and stasis.

## Discussion

Previous studies on mycobacterial phospho-regulation suggest that PstP could play a critical role in modulating cell wall metabolism in the transition between growth and stasis (18, 22, 23, 35, 42, 55, 58, 89). In this work, we explored how the phosphorylation of PstP contributes to this regulation. We report here that the phospho-site T171 of PstP_*Msmeg*_ impacts growth, cell wall metabolism and antibiotic tolerance. We found that the PG master regulator CwlM_*Mtb*_ is a substrate of PstP_*Mtb*_. Our findings indicate that the phosphorylation on PstP affects PG metabolism in stasis and the mycolic acid metabolism during growth.

PG is regulated by phosphorylation factors at several points along the biosynthesis pathway (23, 41, 67, 90), mostly by PknB. PknB’s kinase activity is responsive to lipid II that it detects in the periplasm (91). PstP is a global negative regulator of STPK phosphorylation (42) and has been proposed to be the cognate phosphatase of PknB in regulating cell growth (22, 42, 58, 92). Our data suggest that mutations at T171 of PstP do not affect PG metabolism in growth (Fig. 4A), but that the PstP_*Msmeg*_ T171E strain fails to downregulate PG in starvation (Fig. 4C). We expect that PstP’s activity against the PG regulator CwlM (Fig 2A, top panel) should be critical for this downregulation because it should de-activate MurA, the first enzyme in PG precursor synthesis (23, 40).

The *in vitro* biochemistry (Fig. 2A and B) predicts the log. phase staining data (Fig. 4A), where the *pstP* T171E variant has no difference in apparent PG activity. The proximity of a phospho-site to the substrate binding site of an enzyme may affect the catalytic activity directly (93) but T174 maps to the β-sheet core (β8) in PstP_*Mtb*_, which is distant from the active site (Fig. 1A) (54). The PG staining in starvation suggests that the PstP_*Msmeg*_ T171E phospho-mimetic variant might dephosphorylate CwlM more slowly *in vivo* (Fig. 5B), but this is not what we see *in vitro* (Fig. 2A and B). Therefore, it is possible that, in starvation, phosphorylation at this site may affect PstP’s interaction (94) with other regulatory proteins (95-97) that could modulate PstP’s activity against PG substrates, or it could affect access to substrates via localization changes.

Synthesis of the various mycobacterial cell wall layers are likely synchronized (22, 98). PknB almost surely plays a crucial role in connecting PG and mycolic acid metabolism during growth. If PG metabolism is slowed, PknB could sense the accumulation of periplasmic lipid II (91) and signal to halt mycolic acid biosynthesis by inactivating the FAS-II enzymes and the trehalose monomycolate transporter MmpL3 (92) via phosphorylation (35-37, 59-61, 92). Our data imply that PstP helps balance the effects of these inhibitory phosphorylations to allow coordinated synthesis of mycolic acids in log. phase (Fig. 4B and 5E). Mis-phosphorylation of PstP likely disrupts this coordination and seems to decrease mycolic acid layer metabolism. This may partly explain the slow growth of the *pstP* T171E mutants (Fig. 1B).

We propose that PstP’s regulation of mycolic acid layer biosynthesis occurs in the cytoplasm. DMN-Tre incorporation into the mycomembrane is directly catalyzed by secreted Ag85 enzymes (75, 99), but it is indirectly affected by both cytoplasmic changes in mycolic acid synthesis (75) and altered mycomembrane hydrophobicity (100). PstP and all the STPKs work in the cytoplasm, and there are currently no known systems whereby secreted proteins like Ag85 can be regulated by phosphorylation. All the enzymes of the FAS-II complex, which elongates fatty acids into the long lipids used in mycolic acids (84), are downregulated by phosphorylation (35-37, 59-61), and two are biochemically verified substrates of PstP (35). MmpL3, the mycolic acid flippase (101), is also inhibited by phosphorylation (92). It is likely that PstP could affect the activity of the entire FAS-II complex, including the target of isoniazid, InhA, which is inactivated by threonine phosphorylation (37, 60). Isoniazid is a small hydrophilic drug and undergoes active diffusion via the porins (88, 102); therefore, alterations in mycomembrane permeability are not likely to contribute substantially to differences in isoniazid sensitivity. Although our data (Fig. 5E) does not reveal the exact mis-regulated spot in the mycolic acid synthesis and transport pathway, the higher susceptibility of the phosphomimetic strain (Fig. 5E) to isoniazid suggests that this metabolic pathway is affected. Our DMN-tre staining also suggests that there should be a balance of non-phospho and phospho-form of PstP_*Msmeg*_ T171 (Fig. 4B) during growth to regulate mycomembrane biosynthesis.

PstP may dephosphorylate the cell wall substrates directly, and/ or by de-activating their kinases (103) in both the PG and mycolic acid biosynthesis pathways. All these data combined suggest a complex cross-talk of the STPKs and PstP to regulate diverse cell wall substrates.

## Acknowledgment

This work was supported by grants 1R15GM131317-01 and R01AI148917-01A1 to CCB from the National Institutes of Health, and by startup funds from the University of Texas at Arlington. We thank Kenan Murphy for the plasmids pDE54MCZD and pKM55 and Dirk Schnappinger for pDE43-MCS and RevTetR promoter plasmids used in this study.

---

**Figure S1.**
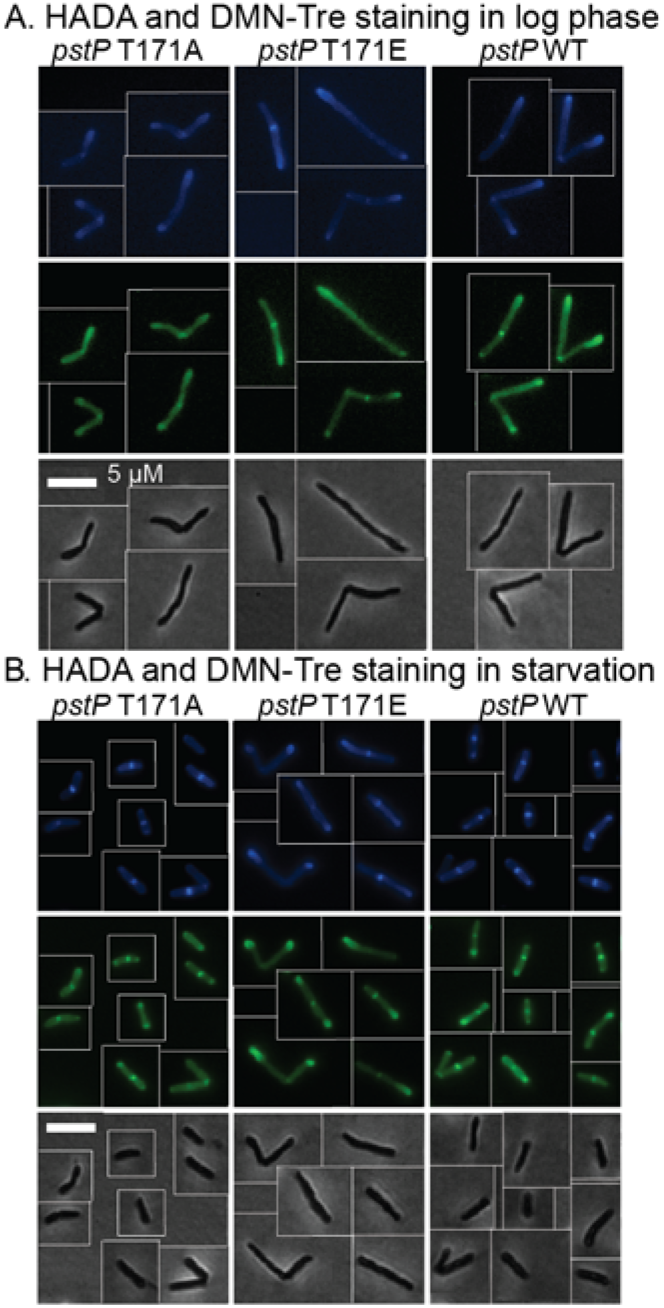
(A) and (B) Representative micrographs of log. phase cells (A) and starved cells in HdB with no glycerol (B) from *pstP* allele strains (WT, T17A and T171E) stained with the fluorescent dyes HADA (blue) and DMN-Tre (green). Corresponding phase images are shown on the bottom panel. The scale bar applies to all images.

**Figure S2:**
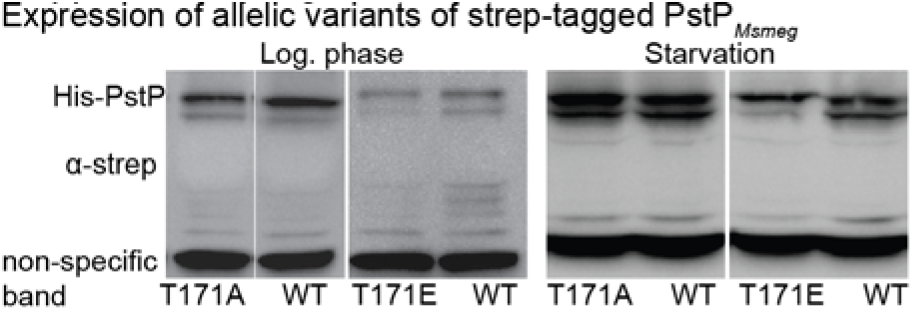
α-strep Western blots of allelic variants of strep-tagged PstP_*Msmeg*_ in log. (left panel) and starvation phase (right panel). A non-specific band at the bottom in all strains was seen in the blots.

